# Genomics-informed insights into microbial degradation of *N,N*-dimethylformamide

**DOI:** 10.1101/2021.03.18.435917

**Authors:** Junhui Li, Paul Dijkstra, Qihong Lu, Shanquan Wang, Shaohua Chen, Deqiang Li, Zhiheng Wang, Zhenglei Jia, Lu Wang, Hojae Shim

## Abstract

Effective degradation of *N,N*-Dimethylformamide (DMF), an important industrial waste product, is challenging as only few bacterial isolates are known to be capable of degrading DMF. Aerobic remediation of DMF has typically been used, whereas anoxic remediation attempts are recently made, using nitrate as one electron acceptor, and ideally include methane as a byproduct. Here, we analyzed 20,762 complete genomes and 28 constructed draft genomes for the genes associated with DMF degradation. We identified 952 genomes that harbor genes involved in DMF degradation, expanding the known diversity of prokaryotes with these metabolic capabilities. Our findings suggest acquisition of DMF-degrading gene via plasmids are important in the order Rhizobiales and genus *Paracoccus*, but not in most other lineages. Degradation pathway analysis reveals that most putative DMF degraders using aerobic Pathway I will accumulate methylamine intermediate, while members of *Paracoccus, Rhodococcus, Achromobacter*, and *Pseudomonas* could potentially mineralize DMF completely under aerobic conditions. The aerobic DMF degradation via Pathway II is more common than thought and is primarily present in α-and β-Proteobacteria and Actinobacteria. Most putative DMF degraders could grow with nitrate anaerobically (Pathway III), however, genes for the use of methyl-CoM to produce methane were not found. These analyses suggest that microbial consortia could be more advantageous in DMF degradation than pure culture, particularly for methane production under the anaerobic condition. The identified genomes and plasmids form an important foundation for optimizing bioremediation of DMF-containing wastewaters.

**Importance:** DMF is extensively used as a solvent in industries, and is classified as a probable carcinogen. DMF is a refractory compound resistant to degradation, and until now, only few bacterial isolates have been reported to degrade DMF. To achieve effective microbial degradation of DMF from wastewater, it is necessary to identify genomic diversity with the potential to degrade DMF and characterize the genes involved in two aerobic degradation pathways and potential anaerobic degradation for methane production. A wide diversity of organisms has the potential to degrade DMF. Plasmid-mediated degradation of DMF is important for Rhizobiales and *Paracoccus*. Most DMF degraders could grow anaerobically with nitrate as electron acceptor, while co-cultures are required to complete intermediate methanogenesis for methane production. This is the first genomics-based global investigation into DMF degradation pathways. The genomic database generated by this study provides an important foundation for the bioremediation of DMF in industrial waste waters.

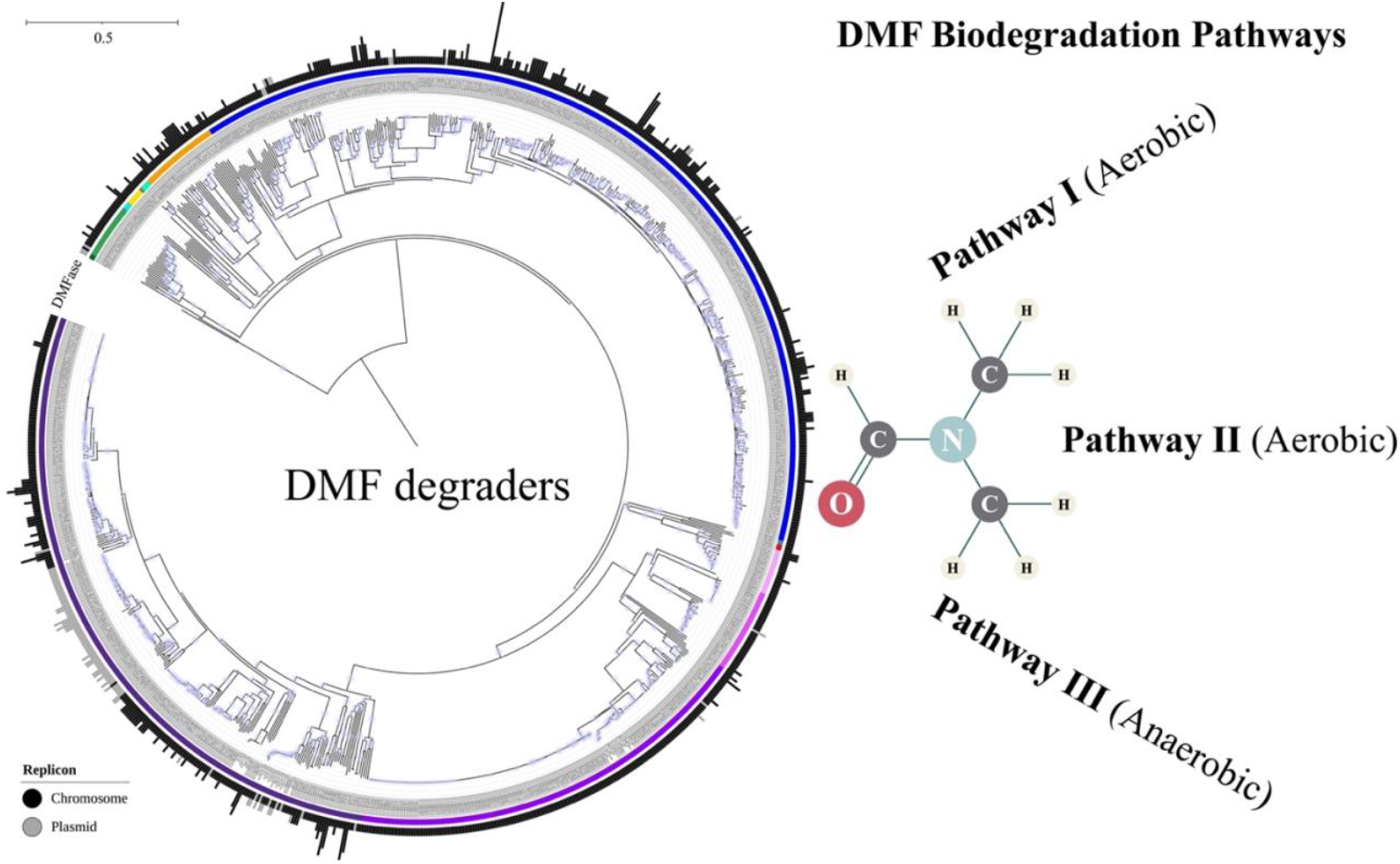

## 1. Introduction

*N,N*-Dimethylformamide (DMF) is a synthetic solvent with low evaporation rate, complete miscibility with water and the majority of other solvents, and is extensively used in synthetic textile, leather, electronics, pharmaceutical and pesticides industries (1). Its use creates industrial wastewater with high levels of DMF, risking contamination of the environment. DMF was classified as probably carcinogenic to humans and is a leading cause of liver disease in chronically exposed workers (2).

Although various physicochemical methods have been used to remove DMF from wastewater (3, 4), microbial degradation is considered to be a superior alternative as it is economical, eco-friendly, and highly efficient (5-9). So far, only a limited number of bacterial isolates have been discovered that are capable of utilizing DMF as the sole carbon and nitrogen source (Table 1). All isolated DMF degraders are aerobic, and mostly from the phylum Proteobacteria, particularly Alphaproteobacteria and Gamaproteobacteria, including members of *Paracoccus, Methylobacterium, Mesorhizobium, Ochrobactrum, Alcaligenes*, and *Pseudomonas*, as well as *Mycobacterium* (Actinobacteria) and *Bacillus* (Firmicutes). Of these bacterial isolates, some (e.g., *Paracoccus* sp. DMF-3, *Alcaligenes* sp. KUFA-1, and *Pseudomonas* sp. DMF 5/8) can grow on high concentration (∼ 50 g/L) DMF solutions (Table 1). Besides *Paracoccus* and *Bacillus*, members of six additional genera (i.e., *Hyphomicrobium, Nitratireductor, Burkholderia, Rhodobacter, Catellibacterium*, and *Bradyrhizobium*) based on marker gene analyses were recently reported as potential DMF degraders under anaerobic conditions (10-12). In our recent study, *Paracoccus* and *Hyphomicrobium* are likely major DMF degraders under aerobic conditions identified using amplicon-based marker gene analyses (13), and supported by identification of genes encoding *N,N*-dimethylformamidase (DMFase) (14). In addition, members of six other genera, i.e., *Achromobacter, Methyloversatilis, Nitratireductor, Pontibaca, Rhodopseudomonas*, and *Starkeya*, carry genes encoding the large and/or small subunits of DMFase (14).

**Table 1.** Summary of properties of bacterial DMF degraders from the literature

| (Sub)Phylum | Genus | Strain | Aerobic | Anaerobic | Nitrate | Max | NMF | DMA | MA | FA | HCHO | Formate | Reference |
| --- | --- | --- | --- | --- | --- | --- | --- | --- | --- | --- | --- | --- | --- |
| α-Proteobacteria | *Paracoccus* | *P. aminophilus* DM-15<br>(= JCM 7686)<br>(GCA_000444995.1) | + | + | + | 30 | + | + | + | + | + | -/+ | (24, 26-30) |
|  |  | *P. aminovorans* DM-82<br>(= JCM 7685)<br>(GCA_900005615.1) | + | NA | + | 30 | + | + | + | + | NA | - | (26, 27) |
|  |  | *P. denitrificans* SD1 | + | + | + | > 28.3 | NA | NA | NA | NA | NA | NA | (31-33) |
|  |  | *Paracoccus* sp. MKU1 | + | NA | + | > 9.4 | NA | NA | NA | NA | NA | NA | (34) |
|  |  | *Paracoccus* sp. MKU2 | + | NA | + | > 9.4 | NA | NA | NA | NA | NA | NA | (34) |
|  |  | *Paracoccus* sp. DMF | + | NA | + | 15 | NA | + | + | + | NA | + | (7) |
|  |  | *Paracoccus* sp. DMF-3 | + | NA | NA | 50 | + | + | + | + | NA | NA | (6) |
|  | *Methylobacterium* | *Methylobacterium* sp. TH-15 | + | NA | NA | 20 | + | + | + | + | NA | - | (27) |
|  |  | *Methylobacterium* sp. DM1<br>(GCA_003111705.1) | + | NA | G | > 2 | NA | + | + | + | + | + | (25) |
|  | *Mesorhizobium* | *M. tamadayense* MM3441 | + | + | + | 40 | + | + | + | + | NA | + | (9) |
|  | *Ochrobactrum* | *Ochrobactrum* sp. DGVK1 | + | + | NA | > 18.9 | NA | + | + | + | NA | + | (8, 35) |
| β-Proteobacteria | *Alcaligenes* | *Alcaligenes* sp. KUFA-1 | + | NA | + | 50 | NA | NA | NA | - | NA | NA | (16, 36) |
| γ-Proteobacteria | *Pseudomonas* | *Pseudomonas* sp. DMF 3/3 | + | NA | + | > 10 | + | + | + | + | NA | + | (16) |
|  |  | *Pseudomonas* sp. DMF 3/4 | + | NA | + | > 10 | + | + | + | + | NA | + | (16) |
|  |  | *Pseudomonas* sp. DMF 3/5 | + | NA | + | > 10 | + | + | + | + | NA | + | (16) |
|  |  | *Pseudomonas* sp. DMF 3/6 | + | NA | + | > 10 | + | + | + | + | NA | + | (16) |
|  |  | *Pseudomonas* sp. DMF 3/11 | + | NA | + | > 10 | + | + | + | + | NA | + | (16) |
|  |  | *Pseudomonas* sp. DMF 3/12 | + | NA | + | > 10 | + | + | + | + | NA | + | (16) |
|  |  | *Pseudomonas* sp. DMF/HW1-5 | + | NA | + | > 10 | + | + | + | + | NA | + | (16) |
|  |  | *Pseudomonas* sp. DMF 4/4 | + | NA | + | > 10 | + | + | + | + | NA | + | (16) |
|  |  | *Pseudomonas* sp. DMF 5/3 | + | NA | + | > 10 | + | + | + | + | NA | + | (16) |
|  |  | *Pseudomonas* sp. DMF 5/5 | + | NA | + | > 10 | + | + | + | + | NA | + | (16) |
|  |  | *Pseudomonas* sp. DMF 5/7 | + | NA | + | > 10 | + | + | + | + | NA | + | (16) |
|  |  | *Pseudomonas* sp. DMF 5/8 | + | NA | + | 56.7 | + | + | + | + | NA | + | (16) |
|  |  | *Pseudomonas* sp. DMF 5/9 | + | NA | + | > 10 | + | + | + | + | NA | + | (16) |
|  |  | *Pseudomonas* sp. DMF 5/10 | + | NA | + | > 10 | + | + | + | + | NA | + | (16) |
|  |  | *Pseudomonas* sp. MBYD-1 | + | NA | NA | > 0.5 | NA | + | + | + | NA | + | (37) |
| Actinobacteria | *Mycobacterium* | *Mycobacterium* sp. TH-35 | + | NA | NA | 30 | + | + | + | - | NA | - | (27) |
| Firmicutes | *Bacillus* | *B. cereus* D-1 | + | + | + | NA | NA | NA | NA | NA | NA | NA | (38) |
|  |  | *B. subtilis* | + | NA | - | NA | NA | NA | NA | NA | NA | NA | (39) |

Max, the maximum concentration of DMF that permits bacterial growth of the isolate (g/L). NMF, *N*-methylformamide; DMA, dimethylamine; MA, methylamine; FA, formamide; HCHO, formaldehyde.
Growth activity: +, positive; -, negative; NA, not available. G, presence of the putative gene. -/+, conflicting results between studies.

Aerobic degradation of DMF is considerably more efficient than anaerobic degradation (11, 15). Therefore, most of the studies have been centered on the aerobic degradation of DMF. Two pathways have been reported for aerobic DMF degradation (16): I) DMFase hydrolyzing DMF to formate and dimethylamine (DMA), followed by converting DMA to methylamine (MA) by dimethylamine dehydrogenase (Fig. 1). This is the most common pathway. II) DMF is degraded via sequential oxidative demethylations, giving rise to *N*-methylformamide (NMF), formaldehyde (HCHO), and formamide (FA), which can be further converted to ammonia and formate by formamidase (Fig. 1). Under aerobic conditions, DMF is ultimately degraded to NH_4_ ^+^ and CO_2_ in both pathways. Anaerobic degradation of DMF by microbial consortia is attracting increasing interest because of the simultaneous energy production (10, 17-20). Pathway III was recently proposed for anaerobic conditions (17). The anaerobic degradation of DMF depends on facultative DMF-degrading bacteria and the presence of intermediate utilizing methanogens for methane production, and requires nitrate as electron acceptor (10, 18). Among the intermediates of aerobic Pathway I, DMA and MA are common substrates for methylotrophic methanogens (21), while formate can be subsequently fermented to methane by methanogens (22). Anaerobic degradation of DMF is possible for denitrifiers utilizing nitrate as an electron acceptor (10). It would be particularly beneficial under anaerobic conditions if DMF degraders are methanogens, which could produce energy as a byproduct of the remediation process. However, it is unclear if such denitrifying and/or methanogenic DMF degraders exist in nature.

**Fig. 1.**
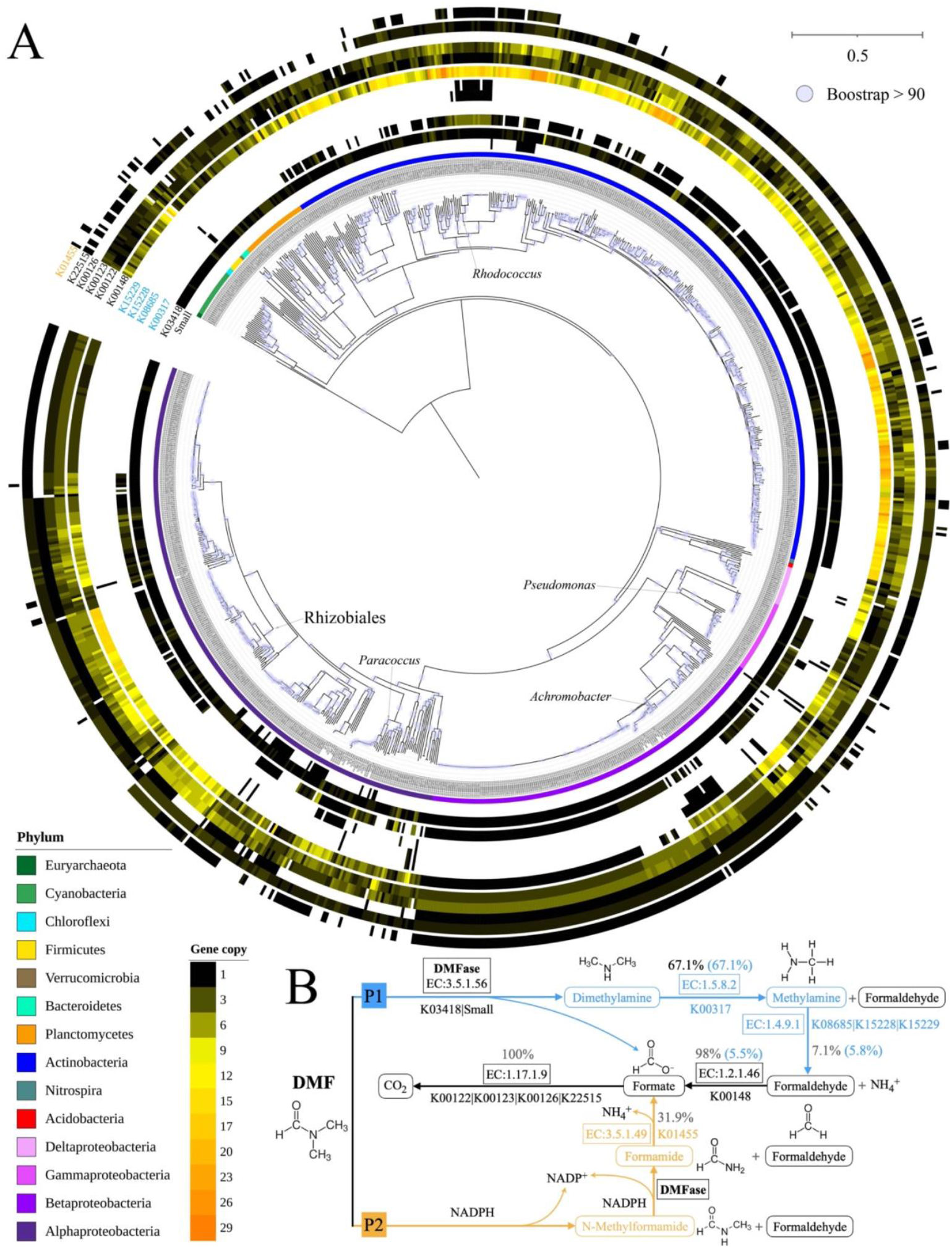
Phylogenetic distribution of putative DMF degraders. A) genes involved in DMF mineralization. The consensus phylogenetic tree was based on 40 single copy marker genes (see Methods for details); B) DMF mineralization pathways under aerobic conditions (34). Enzyme involved in the first step of Pathway II is unknown. Squares indicate enzyme; rounded rectangles indicate produced intermediates. Percentage in black indicates genomes involved in the specific step of the degradation process; percentage in light blue in the brackets indicates genomes involved in each step from DMF to the specific intermediate of the degradation process. Line in light blue indicates pathway I; line in orange indicate pathway II; line in black indicates shared metabolic reactions.

Genes related to DMF degradation are either present on chromosomes or plasmids of bacteria. Multiple strains, e.g., *Paracoccus aminovorans* JCM 7685 (23), *Paracoccus aminophilus* JCM 7686 (24), and *Methylobacterium* sp. DM1 (25), are reported to carry genes encoding DMFase on the plasmid. Likewise, we found that, of 13 *Paracoccus* metagenomic assembled genomes (MAGs) which harbor the large subunit of DMFase, 11 have this gene on their plasmids and 4 contain this gene on the plasmid solely (14). The presence of the gene on plasmids, either as an ancient or recent event, suggests the possibility of horizontal gene acquisition.

In most cases, DMF degrader isolates described in the literature are taxonomically identified using 16S rRNA gene, while complete genomes of most isolates are missing. Here, we analyzed 20,762 complete prokaryotic genomes deposited in GenBank and the 28 MAGs from enriched microbial consortia with DMF as the carrier solvent (13, 14) to 1) identify the putative DMF degraders and metabolic pathways for DMF degradation, 2) test whether the plasmid-mediated degradation of DMF is common, and 3) determine whether the putative DMF degraders have the additional capability to utilize nitrate as an electron acceptor and to break down intermediates of DMF degradation to methane. Overall, the current study substantially expands our knowledge of the diversity of DMF degraders and their potential to function under aerobic and anaerobic conditions.

## 2. Materials and methods

### 2.1. Bioinformatic analyses

To identify putative DMF degraders, the nucleotide sequences of 20,762 complete prokaryotic genomes deposited in GenBank (accessed 11-22-2020) were downloaded and processed using Prodigal (40) to call open reading frames (ORFs). The resulting ORFs were searched against the metabolic pathways of DMF degradation (16) in the KOfam database (41) using hmmscan (E-value < 1e-15 and covered fraction of hmm > 50%) in HMMER V3.2.1 (42). Subsequently, the predicted ORFs of selected genomes harboring at least one copy of the gene for the large and/or small subunit of DMFase were searched against the targeted KO families mapped to pathways of nitrogen (map00910) and methane (map00680) metabolism from the KOfam database (41). The 28 MAGs carrying genes encoding DMFase from enriched microbial consortia (14) were analyzed using the same procedure. Caution should be taken when interpreting the results. A strict cut-off (i.e., e-value < 1e-15 and a coverage > 50% of the gene) was applied to identify the gene in the genomes/MAGs, which may result in the drop of genes. In addition, the presence of a gene is not evidence for the occurrence of the process.

The number of replicons (i.e., chromosome and plasmid) contained in each complete genome were obtained from prokaryotic genome reports (ftp://ftp.ncbi.nlm.nih.gov/genomes/GENOME_REPORTS/prokaryotes.txt). To determine the location of the genes encoding DMFase, we mapped the GenBank accession number to chromosome or plasmid. The location of genes encoding DMFase of the MAGs were based on the predicted plasmid information using Platon V1.2.0 (43) in a separated study (14), whereas the number of replicons contained in the MAGs is unknown.

### 2.2. Phylogenetic visualization

The phylogenetic trees of 952 putative DMF degraders were constructed using Geneious (https://www.geneious.com) with 40 single-copy marker genes (44) which were extracted from the ORFs with fetchMGs v1.2 (45), aligned with Clustalo (46), and trimmed with trimAl v1.2 (-gt 0.5) (47). Likewise, a neighbor-joining phylogenetic tree of 1,243 amino acid sequences of K03418 gene encoding the large submit of DMFase in 952 putative DMF degraders was constructed using Geneious with the same extraction, alignment, and trimming methods using amino acid sequences. Interactive Tree Of Life (iTOL) (48) was applied to visualize the phylogenetic trees.

### 2.3. Data availability

The 28 MAGs containing genes encoding DMFase are available at https://bitbucket.org/junhuilinau/hmm/src/master/.

## 3. Results

### 3.1. Putative N,N-dimethylformamidase in prokaryotes

Among the 20,762 complete prokaryotic genomes, 924 (4.5%) harbor at least one copy of the gene encoding the large and/or small submits of DMFase (Fig. 1a). These genomes are phylogenetically distributed across 11 phyla, including Acidobacteria, Bacteroidetes, Chloroflexi, Cyanobacteria, Firmicutes, Nitrospira, Planctomycetes, and Verrucomicrobia, as well as archaeal Euryarchaeota. This is the first evidence that archaea have the potential for DMF degradation. The majority of genomes are from the Proteobacteria (472) and Actinobacteria (377) phyla. Additionally, 28 MAGs, discovered in metagenomes from 1-year enriched microbial consortia in the presence of DMF as the carrier solvent (13, 14), contained genes encoding DMFase. A single copy of the gene encoding the small submit of DMFase is identified in 41 genomes/MAGs from Alphaproteobacteria (31), the *Mycolicibacterium* genus in Actinobacteria (10), and Betaproteobacteria (1). These results suggest the DMF degraders are far more widely distributed than previously known and include archaea.

The median copy number of the large subunit of DMFase (K03418) is 1, with *Catenulispora acidiphila* DSM 44928 (GCA_000024025.1) within the Actinobacteria phylum harboring the maximum copies of K03418 gene (9). Totally, 216 genomes/MAGs (22.7%) carry multiple copies of K03418 gene, whereas the gene sequences within the same genome could be more similar to that of other genomes. For example, 4 K03418 genes on the chromosome of *Mesorhizobium terrae* (GCA_008727715.1) are distributed in three clades (I.2, III.2, and IV.2), and nine K03418 genes on the chromosome of *Catenulispora acidiphila* DSM 44928 are distributed in clades I.2 and II.7 (Fig. 2).

**Fig. 2.**
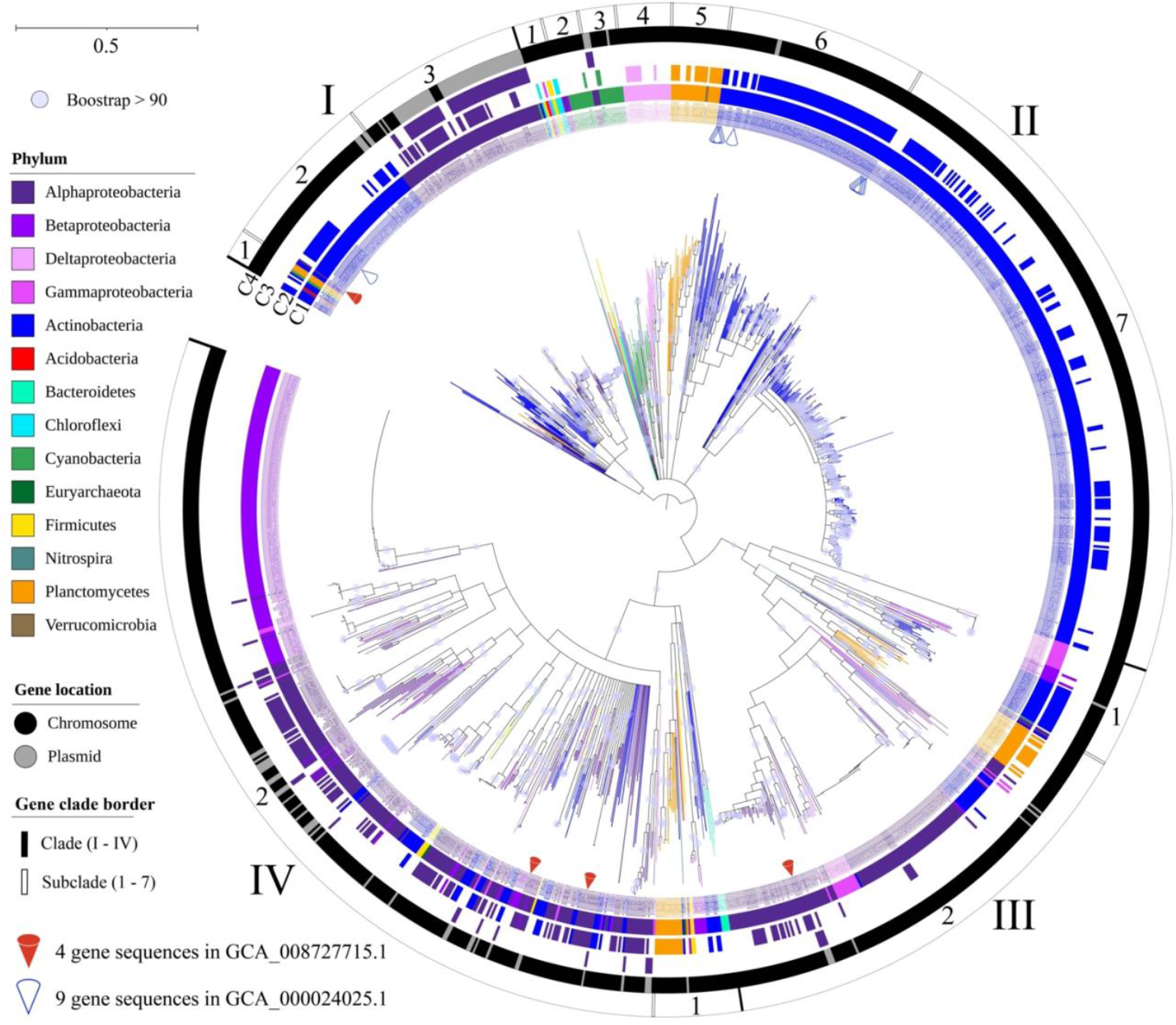
Neighbor-joining phylogenetic tree of 1,243 amino acid sequences of K03418 in 952 putative DMF degraders. The inner circle C1 indicates individual K03418 sequences; the middle circle C2 indicates K03418 sequences that are present in a genome/MAG in multiple copies; the middle circle C3 indicates K03418 sequences in the order of Rhizobiales; the outer circle C4 indicates gene location (chromosome or plasmid). Colors of inner circle (C1), middle circles (C2, C3), and the labels indicate phyla.

Next, we assessed the potential for degradation of intermediates in the DMF degradation pathways under the aerobic conditions (Fig. 1). In the case of Pathway I, 67.1% (639/952) of genomes/MAGs harbor K00317 encoding dimethylamine dehydrogenase [EC:1.5.8.2]. These genomes/MAGs belong to Firmicutes, Actinobacteria, and α-, β-, γ-Proteobacteria (Fig. 1a). Only 5.8% (55/952) genomes/MAGs carry at least one copy of genes (K08685, K15228, or K15229) encoding MA dehydrogenase [EC:1.4.9.1]. Thirteen additional genomes/MAGs carry genes encoding MA dehydrogenase but none for DMA dehydrogenase. Of the 55 putative DMA and MA degraders, 3 genomes don ‘t carry K00148 encoding HCHO dehydrogenase [EC:1.2.1.46] which converts HCHO to F, although K00148 is present in 98% (933/952) of the DMF degrading genomes/MAGs (median copy 8). Moreover, all 68 putative MA degraders carry K00148.

In Pathway II, K01455 encoding formamidase [EC:3.5.1.49] converts FA to formate and is present in 304 genomes/MAGs. The gene involved in formate oxidation is ubiquitous, i.e., at least one of the genes (K00122, K00123, K00126, and K22515) encoding formate dehydrogenase is present in all 952 genomes/MAGs. MA is often the end-product of the DMF degradation (Pathway I) in most putative DMF degraders, as the genes encoding MA dehydrogenase [EC:1.4.9.1] are absent. For 52 genomes/MAGs, genes for complete mineralization of DMF to CO_2_ are present, including 25 genomes/MAGs that carry the formamidase gene (including 15 *Rhodococcus* (Actinobacteria) and 7 *Paracoccus* MAGs (α-Proteobacteria)).

Most DMF degrader isolates from literature (Table 1) are capable of degrading NMF (20 isolates), DMA (24) and MA (24), whereas 23 tested positive and 2 negative for FA degradation activity, and 19 tested positive and 3 negative for formate degradation activity (Table 1). Results for *Paracoccus aminophilus* DM-15 (= JCM 7686) were contradictory (26, 30). Our genomic analysis supports the idea that this isolate is capable of oxidizing formate (Fig. 1).

### 3.2. Plasmid-mediated degradation of DMF

It is estimated that 1.6-32.6% of the genes of each prokaryotic genome has been acquired via horizontal gene transfer (HGT) (49). Previous studies have provided evidence of plasmid-borne genes encoding DMFase (14, 24, 25). However, it is unclear whether this gene is contained in plasmids frequently because of the low number of sequenced genomes of DMF degraders. Out of the 952 genomes/MAGs, 34.2% (326) carry at least one plasmid, whereas 90% (857) carry one or more copies of K03418 only on the chromosome. A total of 95 genomes/MAGs, accounting for 29.1% of 326 genomes with plasmid, harbor the K03418 gene on the plasmid, while 18 genomes carry the gene on both the chromosome and the plasmid and 77 genomes harbor K03418 only on the plasmid. Notably, all 59 putative DMF degraders in the Rhizobiales order of α-Proteobacteria only harbor K03418 on their plasmids, with the exception of *Ensifer mexicanus* ITTG R7 (GCA_013488225.1) and *Ciceribacter thiooxidans* F43B (GCA_014126615.1) which harbor K03418 on both the plasmid and chromosome (Fig. 3). Multiple copies (median 4) of plasmids in Rhizobiales genomes could enhance the opportunity to acquire K03418 gene. Our analyses suggest that plasmids play an important role in acquisition of genes encoding DMFase in the order Rhizobiales.

**Fig. 3.**
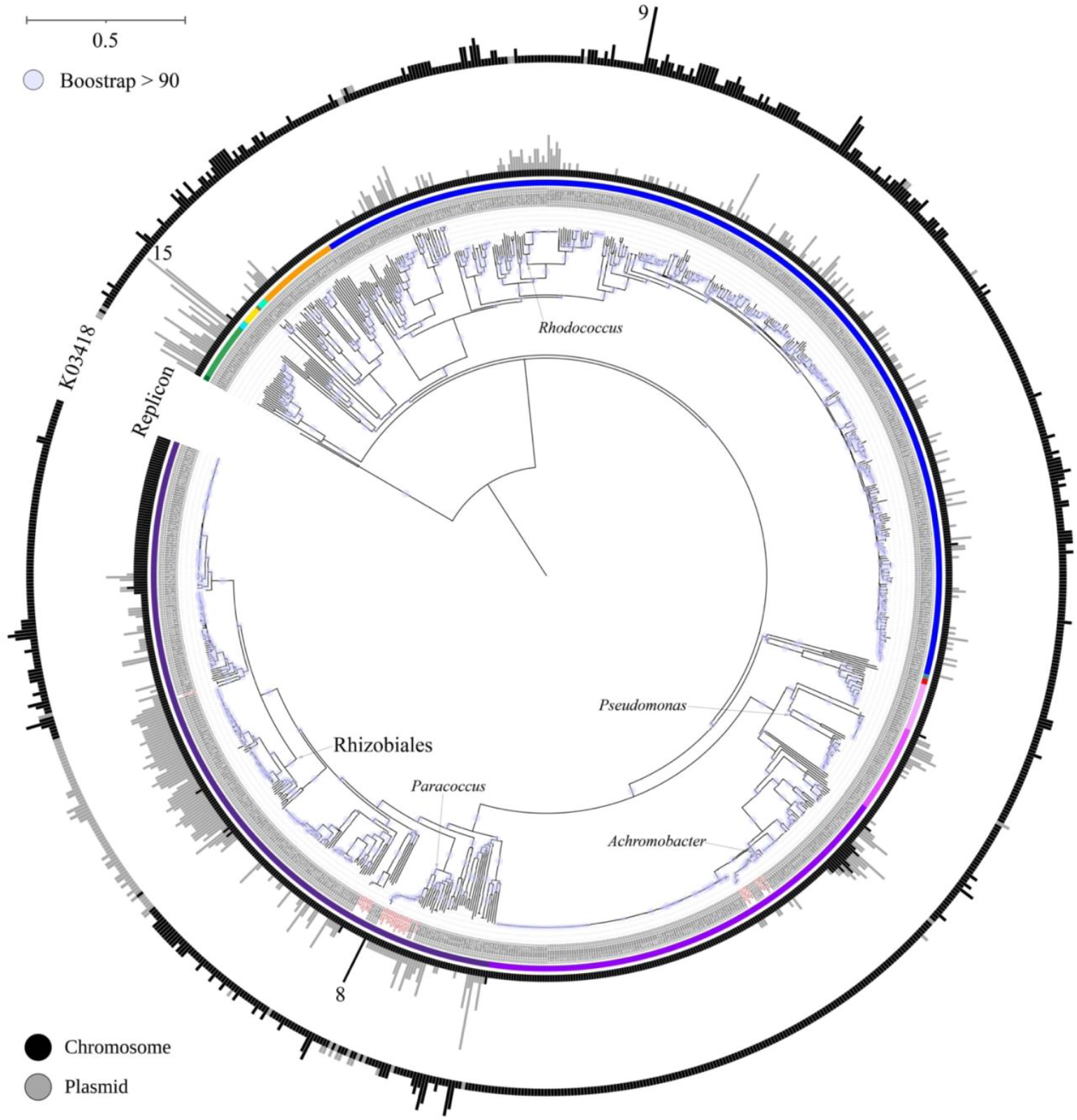
Presence of DMFase (K03418) on plasmids. The inner bar indicates the copy number of replicons (chromosome and plasmid), the outer bar indicates the copy number of K03418. The MAGs in red indicate the presence or absence of plasmids estimated in a separate study (27) rather than the specific copies of chromosome and plasmid.

The majority of K03418 sequences (60) in Rhizobiales genomes clustered together in clade I.3 with other α-Proteobacteria genomes, while the remaining 18 K03418 sequences in Rhizobiales genomes are distributed in three other clades (i.e., II.3, III.2, and IV.2). Among which, II.3 consists of 3 Rhizobiales and 12 Cyanobacteria genomes; III.2 consists of 4 Rhizobiales with similar K03418 sequences as other α-Proteobacteria; and IV.2 consists of 11 Rhizobiales genomes, with similar K03418 sequences to that of Actinobacteria and other α-, β-, γ-Proteobacteria (Fig. 2). Likewise, multiple *Paracoccus* genomes carry K03418 only on the plasmid. These results suggest that plasmids play a critical role in DMF degradation, particularly for Rhizobiales and *Paracoccus*.

In addition, both putative archaeal DMF degraders, *Haloferax gibbonsii* LR2 and *Halalkalicoccus jeotgali* B3, carry a single copy of K03418 on their plasmids. Nineteen out of 23 Cyanobacteria genomes carry plasmids (median 5 copies), including *Nostoc* sp. C057 (GCA_013393925.1) which carries the most copies (15) of plasmids among all DMF degraders, whereas only 1 Cyanobacteria genome harbors K03418 on the plasmid. It appears that plasmid-mediated HGT of K03418 is rare in Cyanobacteria, indicative of vertical descent. This is further supported by the evolutionary analyses of K03418 gene, i.e., the K03418 sequences of Cyanobacteria are clustered together in II.2 and II.3 (Fig. 2), in contrast to Proteobacteria, Actinobacteria and Planctomycetes where they are not clustered together.

### 3.3. Anaerobic degradation of DMF with nitrate but no methane production

The anaerobic degradation of DMF, i.e., Pathway III, has been tested in lab-scale anaerobic bioreactors (10, 18). However, culturing the DMF-degrading bacteria under anaerobic conditions is difficult. Although all isolates metabolize DMF aerobically, several are known as facultative anaerobes (Table 1), e.g., *Paracoccus aminophilus* JCM 7686, *Paracoccus denitrificans* SD1, *Mesorhizobium tamadayense* MM3441, *Ochrobactrum* sp. DGVK1, and *Bacillus cereus* D 1. A capability of DMF degraders to conduct denitrification would offer advantage for removal of DMF in environments with low oxygen availability (9). Anaerobic degradation of DMF is possible for denitrifiers, which use nitrate as an electron acceptor, and most of these isolated DMF degraders are capable of reducing nitrate (Table 1).

We analyzed the presence of nitrate reduction genes involved in three pathways, i.e., assimilatory nitrate reduction (to ammonium), dissimilatory nitrate reduction (to ammonium), and denitrification to N_2_. Of 952 putative DMF degraders, 96.4% (918) possess genes for all steps of at least one of the three nitrate reduction pathways (Fig. 4). About 95.2% (906) harbor genes for assimilatory nitrate reduction to ammonium; 40.2% (383) harbor the genes involved in dissimilatory nitrate reduction to ammonium; and 10.8% (103) harbor all genes involved in each step of denitrification. These results suggest that most putative DMF degraders are likely to grow under the anaerobic condition while utilizing nitrate as electron acceptor. The addition of nitrate to anaerobic bioreactors therefore could be sufficient for DMF degradation under anaerobic conditions (5).

**Fig. 4.**
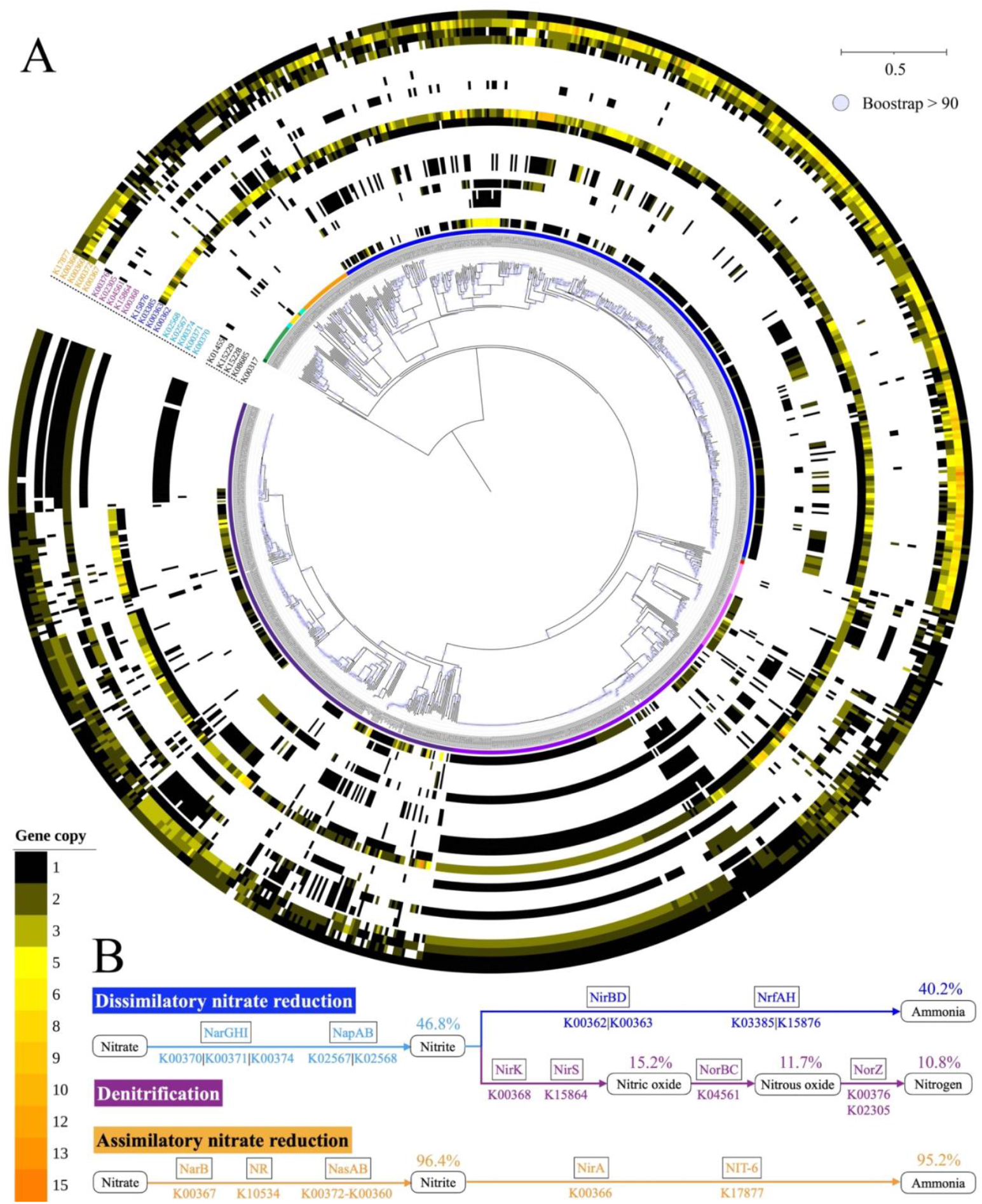
Nitrate reduction of putative DMF degraders. A) Enzymes and genes involved in nitrate reduction; B) three nitrate reduction pathways (KEGG map00910). Squares indicate abbreviated enzyme names; rounded rectangles indicate produced intermediates. Percentage indicates genomes involved in each step from DMF to the specific intermediate of the degradation process. Line in light blue indicates the first step of dissimilatory nitrate reduction and denitrification; line in blue indicates the second step of dissimilatory nitrate reduction; line in purple indicates the other steps of denitrification; line in orange indicates the assimilatory nitrate reduction.

A total of 632 (66.4%) putative DMF degraders are potential denitrifiers that also possess K00317 encoding DMA dehydrogenase, converting DMA to MA and HCHO, and among which 55 genomes/MAGs harbor genes involved in MA oxidation. A total of 616 genomes/MAGs possess K00148 which converts HCHO to formate (Fig. 4a). Additionally, 299 putative denitrifiers, carrying K01455, have the potential to convert FA (Pathway II intermediate) to F. Moreover, all putative denitrifiers can utilize formate as substrate. It appears the putative denitrifiers among the DMF degraders are capable of utilizing the metabolic intermediates, particularly formate and HCHO, to grow under the anaerobic condition.

One advantage of anaerobic DMF degradation is energy production. Simultaneous methane production was achieved while degrading DMF using microbial consortium, albeit maintaining continuous culture is difficult (10, 18). We next analyzed the potential of putative DMF degraders to produce methane (Fig. 5). Although 910 genomes/MAGs possess the genes involved in the conversion of DMA to methyl-CoM, the genes for methyl-CoM reductase [EC:2.8.4.1] needed to produce methane was not observed. In addition, none of the genomes/MAGs carry genes for converting MA to methyl-CoM. These results suggest that microbial consortia are required that include methanogens to complete the methanogenesis of DMA and MA for methane production. Moreover, when formate is used as the electron donor, it is oxidized by formate dehydrogenase [EC:1.17.98.3, EC:1.8.98.6] to CO_2_ and reduced coenzyme F420, which is a required intermediate for methane production (50). Although 929 genomes/MAGs have the potential to oxidize formate to reduced coenzyme F420, none of the genomes harbors the genes encoding methyl-CoM reductase [EC:2.8.4.1] involved in the last step from methyl-CoM to methane.

**Fig. 5.**
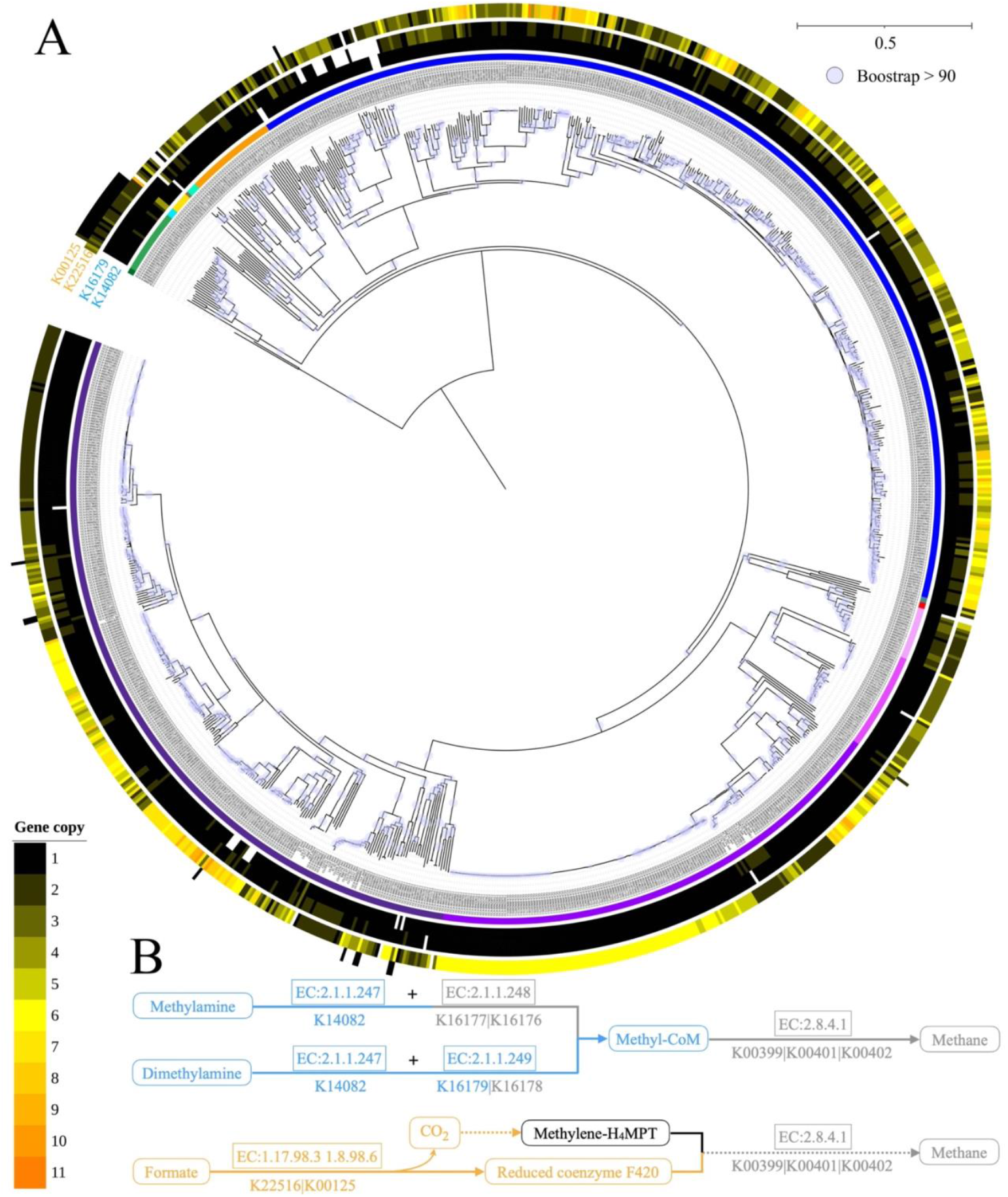
Methanogenesis potential of putative DMF degraders. A) Enzymes and genes involved in the methanogenesis of dimethylamine (DMA), methylamine (MA), and formate (F); B) methanogenesis pathways (DMA and MA: KEGG map00680; formate based on previously established pathways (63). Squares indicate enzyme; rounded rectangles indicate metabolic intermediates. Line in light blue indicates methylotrophic methanogenesis; line in orange indicates hydrogenotrophic methanogenesis; line in grey indicates pathway without identified gene; line in black indicates not tested pathway; dotted line indicates multiple processes.

Together these findings suggest over 50% of the putative DMF degraders could grow anaerobically while utilizing nitrate as electron acceptor and degrade the hydrolyzed intermediates, whereas other methanogens are required to complete the methanogenesis of DMA to produce methane.

## 4. Discussion

DMF is a refractory compound resistant to degradation, and until now, only a small number of bacterial isolates have been reported to degrade DMF under aerobic conditions. However, recent studies using 16S rRNA gene-based identification under both aerobic (13) and anaerobic conditions (10-12) suggest a much wider distribution of DMF degraders.

Our results from the genomic analysis indicate that a total of 4.5% of publicly available full prokaryotic genomes are putative DMF degraders and are far more widely distributed than previously known. Besides genomes in Proteobacteria, Actinobacteria, and Firmicutes phyla, putative DMF degraders are present in 8 other phyla, including two archaeal lineages. Knowledge, isolation and further characterization of these genomes would be particularly useful for developing biodegradation systems, either as pure culture or as enriched microbial consortia. DMF is miscible with water and the majority of organic solvents (1) and has been widely used as carrier solvent in enrichment or isolation of bacteria capable of degrading other water insoluble xenobiotic compounds. In this regard, our results further strengthen our recent statement that when isolating xenobiotic degraders, the presence of the carrier solvent should not be ignored (13). For instance, *Mesorhizobium tamadayense* MM3441 was initially enriched and isolated to degrade pyrene with DMF as the carrier solvent (9), however it is also able to degrade DMF. The 28 MAGs (*Achromobacter* spp., *Hyphomicrobium* spp., *Nitratireductor* sp., and *Paracoccus* spp.) harboring genes encoding DMFase included in this study were also enriched to degrade other xenobiotic compounds in the presence of DMF (14). DMF should be cautiously used as carrier solvent for enriching microbial cultures to degrade other xenobiotic compounds due to it may result in the co-selection of other bacteria which are capable of utilizing DMF as substrates and/or the target xenobiotic compounds (9, 13).

Despite it is unclear when the HGT events occurred, the carried genes encoding DMFase on plasmids plays an important role in DMF degradation. However, the importance of plasmid-mediated gene acquisition differs between taxa. Although the gene encoding DMFase is frequently found in the order of Rhizobiales and the genus *Paracoccus* of α-Proteobacteria, it was relatively rare in Cyanobacteria. Horizontal gene acquisition is important for and common in the order of Rhizobiales ss(51) and recognized as a major driving force in the evolution of lifestyles in this order (52, 53). Likewise, HGT via plasmids is also considered as the driving force of *Paracoccus* evolution (29, 54). Our analyses on plasmid carriage further support the important role of plasmid in gene acquisition occurred in *Paracoccus*, i.e., 19 out of 21 complete *Paracoccus* genomes deposited in GenBank carry plasmid.

Most of the putative DMF degraders may result in the accumulation of MA intermediates for organisms metabolizing DMF via Pathway I because only around 6% of the DMF degraders have the potential to convert MA to HCHO (Fig. 1b). DMF degraders containing genes involved in MA degradation are primarily found in the genera *Paracoccus, Rhodococcus, Achromobacter*, and *Pseudomonas* (Fig. 1). These taxa could potentially mineralize DMF completely to CO_2_ via Pathway I. DMF degraders without genes for MA degradation could still be useful for DMF degradation when used as part of a microbial consortium/ mixed cultures. DMF degradation via Pathway II was thought to be less common (16), whereas the genes encoding the N-demethylase for the degradation of DMF and NMF via oxidative demethylations are not known. Therefore, we are not able at this point to evaluate the genomic diversity involved in the sequential oxidative demethylations of DMF in Pathway II. This knowledge gap requires identification of the N-demethylase genes involved in the sequential oxidative demethylation reactions. Nevertheless, among known bacterial DMF degraders, all tested isolates are capable of utilizing NMF as the sole carbon source (Table 1). Moreover, near 30% of putative DMF degraders carry the gene for the degradation of FA (intermediate of Pathway II), and almost all contain the genes for the degradation of formate and HCHO (intermediates of both Pathways I and II). DMF degraders containing genes for FA degradation are distributed across multiple phyla, but primarily present in α-, β-Proteobacteria and Actinobacteria classes, including *Rhodococcus* and *Paracoccus* genera. Most putative DMF degraders seem to be capable of growing anaerobically with nitrate as electron acceptor. However, Pathway III is incomplete because genes encoding methyl-CoM reductase as the last step of methanogenesis are absent. Individual taxa seem unlikely to complete the DMF methanogenic degradation with methane production, and other methanogens are required to complete the methanogenesis of hydrolyzed intermediates. Thus, microbial consortia have advantages by enabling co-metabolism and complementary enzymes in Pathway III. Recently, it was shown that the effective methanogenic degradation of DMF could not be maintained during the long-term operation of anaerobic membrane (10) and up-flow anaerobic sludge blanket bioreactors (10, 18). Nitrate addition improves the performance of an anoxic denitrification reactor (5). Yet, it is not clear how nitrate would influence methane production. Nitrate dose is expected to play a key role for methanogenic degradation of DMF as heterotrophic denitrification takes over methanogenesis when carbon: nitrogen < 5 (55), highlighting the need for further studies on the effect of nitrate on methanogenic degradation of DMF.

## 5. Conclusions

Taken together, our results reveal that 952 fully sequenced genomes harbor genes encoding DMFase, and are phylogenetically distributed across 11 phyla, substantially expanding the functional diversity in DMF degradation. Plasmid-mediated degradation of DMF plays critical roles in certain taxa, e.g., order Rhizobiales and genus *Paracoccus*. Although many members of *Paracoccus, Rhodococcus, Achromobacter*, and *Pseudomonas* genera appear to be able to potentially mineralize DMF completely via Pathway I under aerobic conditions, mixed microbial cultures probably are needed for DMF degradation particularly via Pathway III, where methanogens are required to complete the methanogenesis of DMF degradation intermediates. This study provides in-depth information on genome-scale metabolic pathways in DMF degradation and their phylogenetic distribution.

## Acknowledgements

This work was supported by the National Natural Science Foundation of China (Grant Nos. 51409106, U1901601) and the University of Macau Multi-Year Research Grant (MYRG2018-00108-FST).

## Reference

1. HSDB. 2015. Hazardous Substance Data Base. Bethesda (MD), USA: United States National Library of Medicine. Available from: http://toxnet.nlm.nih.gov/cgi-bin/sis/htmlgen?HSDB.

2. Kim TH, Kim SG. 2011. Clinical Outcomes of Occupational Exposure to N,N-Dimethylformamide: Perspectives from Experimental Toxicology. Safety and Health at Work 2:97–104.

3. Zhao Y, Ma L, Chang W, Huang Z, Feng X, Qi X, Li Z. 2018. Efficient photocatalytic degradation of gaseous N,N-dimethylformamide in tannery waste gas using doubly open-ended Ag/TiO2 nanotube array membranes. Applied Surface Science 444:610–620.

4. Dou P, Song J, Zhao S, Xu S, Li X, He T. 2019. Novel low cost hybrid extraction-distillation-reverse osmosis process for complete removal of N,N-dimethylformamide from industrial wastewater. Process Safety and Environmental Protection 130:317–325.

5. Wang J, Liu X, Jiang X, Zhang L, Hou C, Su G, Wang L, Mu Y, Shen J. 2020. Facilitated bio-mineralization of N, N-dimethylformamide in anoxic denitrification system: Long-term performance and biological mechanism. Water Research 186:116306.

6. Zhou X, Jin W, Sun C, Gao S-H, Chen C, Wang Q, Han S-F, Tu R, Latif MA, Wang Q. 2018. Microbial degradation of N,N-dimethylformamide by Paracoccus sp. strain DMF-3 from activated sludge. Chemical Engineering Journal 343:324–330.

7. Swaroop S, Sughosh P, Ramanathan G. 2009. Biomineralization of N,N-dimethylformamide by Paracoccus sp. strain DMF. Journal of Hazardous Materials 171:268–272.

8. Veeranagouda Y, Emmanuel Paul PV, Gorla P, Siddavattam D, Karegoudar TB. 2006. Complete mineralisation of dimethylformamide by Ochrobactrum sp. DGVK1 isolated from the soil samples collected from the coalmine leftovers. Applied Microbiology and Biotechnology 71:369–375.

9. Dhar K, Subashchandrabose SR, Venkateswarlu K, Megharaj M. 2020. Mesorhizobium tamadayense MM3441: A novel methylotroph with a great potential in degrading N, N′-dimethylformamide. International Biodeterioration & Biodegradation 153:105045.

10. Li L, Kong Z, Xue Y, Wang T, Kato H, Li Y-Y. 2020. A comparative long-term operation using up-flow anaerobic sludge blanket (UASB) and anaerobic membrane bioreactor (AnMBR) for the upgrading of anaerobic treatment of N, N-dimethylformamide-containing wastewater. Science of The Total Environment 699:134370.

11. Kong Z, Li L, Wu J, Zhang T, Li Y-Y. 2019. Insights into the methanogenic degradation of N, N-dimethylformamide: The functional microorganisms and their ecological relationships. Bioresource Technology 271:37–47.

12. Kong Z, Li L, Kurihara R, Kubota K, Li Y-Y. 2018. Anaerobic treatment of N, N-dimethylformamide-containing wastewater by co-culturing two sources of inoculum. Water Research 139:228–239.

13. Li J, Wu C, Chen S, Lu Q, Shim H, Huang X, Jia C, Wang S. 2020. Enriching indigenous microbial consortia as a promising strategy for xenobiotics ‘ cleanup. Journal of Cleaner Production 261:121234.

14. Li J, Jia C, Lu Q, Hungate BA, Dijkstra P, Wang S, Wu C, Chen S, Li D, Shim H. 2021. Mechanistic insights into the success of xenobiotic degraders resolved from metagenomes of microbial enrichment cultures. bioRxiv doi:10.1101/2021.03.03.433815:2021.03.03.433815.

15. Bromley-Challenor KCA, Caggiano N, Knapp JS. 2000. Bacterial growth on N,N-dimethylformamide: implications for the biotreatment of industrial wastewater. Journal of Industrial Microbiology and Biotechnology 25:8–16.

16. Ghisalba O, Cevey P, Küenzi M, Schär H-P. 1985. Biodegradation of chemical waste by specialized methylotrophs, an alternative to physical methods of waste disposal. Conservation & Recycling 8:47–71.

17. Kong Z, Li L, Kurihara R, Zhang T, Li Y-Y. 2019. Anaerobic treatment of N, N-dimethylformamide-containing high-strength wastewater by submerged anaerobic membrane bioreactor with a co-cultured inoculum. Science of The Total Environment 663:696–708.

18. Kong Z, Li L, Wang T, Rong C, Xue Y, Zhang T, Wu J, Li Y-Y. 2020. New insights into the cultivation of N, N-dimethylformamide-degrading methanogenic consortium: A long-term investigation on the variation of prokaryotic community inoculated with activated sludge. Environmental Research 182:109060.

19. Kong Z, Li L, Li Y-Y. 2019. Long-term performance of UASB in treating N, N-dimethylformamide-containing wastewater with a rapid start-up by inoculating mixed sludge. Science of The Total Environment 648:1141–1150.

20. Kong Z, Li L, Li Y-Y. 2018. Characterization and variation of microbial community structure during the anaerobic treatment of N, N-dimethylformamide-containing wastewater by UASB with artificially mixed consortium. Bioresource Technology 268:434–444.

21. Hippe H, Caspari D, Fiebig K, Gottschalk G. 1979. Utilization of trimethylamine and other N-methyl compounds for growth and methane formation by Methanosarcina barkeri. Proceedings of the National Academy of Sciences of the United States of America 76:494–498.

22. Kurth JM, Op den Camp HJM, Welte CU. 2020. Several ways one goal—methanogenesis from unconventional substrates. Applied Microbiology and Biotechnology 104:6839–6854.

23. Czarnecki J, Dziewit L, Puzyna M, Prochwicz E, Tudek A, Wibberg D, Schlüter A, Pühler A, Bartosik D. 2017. Lifestyle-determining extrachromosomal replicon pAMV1 and its contribution to the carbon metabolism of the methylotrophic bacterium Paracoccus aminovorans JCM 7685. Environmental Microbiology 19:4536–4550.

24. Dziewit L, Dmowski M, Baj J, Bartosik D. 2010. Plasmid pAMI2 of Paracoccus aminophilus JCM 7686 carries N,N-dimethylformamide degradation-related genes whose expression is activated by a LuxR family regulator. Applied and Environmental Microbiology 76:1861.

25. Lu X, Wang W, Zhang L, Hu H, Xu P, Wei T, Tang H. 2019. Molecular Mechanism of N,N-Dimethylformamide Degradation in Methylobacterium sp. Strain DM1. Applied and environmental microbiology 85:e00275–19.

26. Urakami T, Araki H, Oyanagi H, Suzuki K-I, Komagata K. 1990. Paracoccus aminophilus sp. nov. and Paracoccus aminovorans sp. nov., which Uuilize N,N-dimethylformamide. Int J Syst Bacteriol 40:287–291.

27. Urakami T, Kobayashi H, Araki H. 1990. Isolation and identification of N,N-Dimethylformamide-biodegrading bacteria. Journal of Fermentation and Bioengineering 70:45–47.

28. Dziewit L, Kuczkowska K, Adamczuk M, Radlinska M, Bartosik D. 2011. Functional characterization of the type II PamI restriction-modification system derived from plasmid pAMI7 of Paracoccus aminophilus JCM 7686. FEMS Microbiology Letters 324:56–63.

29. Dziewit L, Czarnecki J, Wibberg D, Radlinska M, Mrozek P, Szymczak M, Schlüter A, Pühler A, Bartosik D. 2014. Architecture and functions of a multipartite genome of the methylotrophic bacterium Paracoccus aminophilus JCM 7686, containing primary and secondary chromids. BMC Genomics 15:124.

30. Dziewit L, Czarnecki J, Prochwicz E, Wibberg D, Schlüter A, Pühler A, Bartosik D. 2015. Genome-guided insight into the methylotrophy of Paracoccus aminophilus JCM 7686. Frontiers in Microbiology 6:852.

31. Siddavattam D, Karegoudar TB, Mudde SK, Kumar N, Baddam R, Avasthi TS, Ahmed N. 2011. Genome of a novel isolate of Paracoccus denitrificans capable of degrading N,N-dimethylformamide. Journal of Bacteriology 193:5598.

32. Sanjeevkumar S, Nayak AS, Santoshkumar M, Siddavattam D, Karegoudar TB. 2013. Paracoccus denitrificans SD1 mediated augmentation with indigenous mixed cultures for enhanced removal of N,N-dimethylformamide from industrial effluents. Biochemical Engineering Journal 79:1–6.

33. Zheng Y, Chen D, Li N, Xu Q, Li H, He J, Lu J. 2016. Efficient simultaneous adsorption-biodegradation of high-concentrated N,N-dimethylformamide from water by Paracoccus denitrificans-graphene oxide microcomposites. Scientific Reports 6:20003.

34. Nisha KN, Devi V, Varalakshmi P, Ashokkumar B. 2015. Biodegradation and utilization of dimethylformamide by biofilm forming Paracoccus sp. strains MKU1 and MKU2. Bioresource Technology 188:9–13.

35. Kumar SS, Kumar MS, Siddavattam D, Karegoudar T. 2012. Generation of continuous packed bed reactor with PVA–alginate blend immobilized Ochrobactrum sp. DGVK1 cells for effective removal of N, N-dimethylformamide from industrial effluents. Journal of hazardous materials 199:58–63.

36. Hasegawa Y, Matsuo M, Sigemoto Y, Sakai T, Tokuyama T. 1997. Purification and characterization of N, N-dimethylformamidase from Alcaligenes sp. KUFA-1. Journal of fermentation and bioengineering 84:543–547.

37. Yang S, Chen L, Gu H, Cai T. 2011. Isolation, identification and degradation characteristics of a DMF-degrading bacterial MBYD-1. China Environmental Science 31:437–442.

38. Okazaki M, Hamada T, Fujii H, Kusudo O, Mizobe A, Matsuzawa S. 1995. Development of poly(vinyl alcohol) hydrogel for waste water cleaning. II. Treatment of N,N-dimethylformamide in waste water with poly(vinyl alcohol) gel with immobilized microorganisms. Journal of Applied Polymer Science 58:2243–2249.

39. Vidhya R, Thatheyus A. 2013. Biodegradation of dimethylformamide using Bacillus subtilis. Am J Microbiol Res 1:10–15.

40. Hyatt D, Chen G-L, Locascio PF, Land ML, Larimer FW, Hauser LJ. 2010. Prodigal: prokaryotic gene recognition and translation initiation site identification. BMC bioinformatics 11:119–119.

41. Aramaki T, Blanc-Mathieu R, Endo H, Ohkubo K, Kanehisa M, Goto S, Ogata H. 2020. KofamKOALA: KEGG Ortholog assignment based on profile HMM and adaptive score threshold. Bioinformatics 36:2251–2252.

42. Finn RD, Clements J, Eddy SR. 2011. HMMER web server: interactive sequence similarity searching. Nucleic Acids Research 39:W29–W37.

43. Schwengers O, Barth P, Falgenhauer L, Hain T, Chakraborty T, Goesmann A. 2020. Platon: identification and characterization of bacterial plasmid contigs in short-read draft assemblies exploiting protein-sequence-based replicon distribution scores. bioRxiv doi:10.1101/2020.04.21.053082:2020.04.21.053082.

44. Wu D, Jospin G, Eisen JA. 2013. Systematic identification of gene families for use as “markers “ for phylogenetic and phylogeny-driven ecological studies of bacteria and archaea and their major subgroups. PLOS ONE 8:e77033.

45. Mende DR, Sunagawa S, Zeller G, Bork P. 2013. Accurate and universal delineation of prokaryotic species. Nature Methods 10:881–884.

46. Sievers F, Wilm A, Dineen D, Gibson TJ, Karplus K, Li W, Lopez R, McWilliam H, Remmert M, Söding J, Thompson JD, Higgins DG. 2011. Fast, scalable generation of high-quality protein multiple sequence alignments using Clustal Omega. Molecular Systems Biology 7:539–539.

47. Capella-Gutiérrez S, Silla-Martínez JM, Gabaldón T. 2009. trimAl: a tool for automated alignment trimming in large-scale phylogenetic analyses. Bioinformatics (Oxford, England) 25:1972–1973.

48. Letunic I, Bork P. 2016. Interactive tree of life (iTOL) v3: an online tool for the display and annotation of phylogenetic and other trees. Nucleic Acids Research 44:W242–W245.

49. Koonin EV, Makarova KS, Aravind L. 2001. Horizontal Gene Transfer in Prokaryotes: Quantification and Classification. Annual Review of Microbiology 55:709–742.

50. Costa KC, Wong PM, Wang T, Lie TJ, Dodsworth JA, Swanson I, Burn JA, Hackett M, Leigh JA. 2010. Protein complexing in a methanogen suggests electron bifurcation and electron delivery from formate to heterodisulfide reductase. Proceedings of the National Academy of Sciences 107:11050.

51. Wang S, Meade A, Lam H-M, Luo H. 2020. Evolutionary Timeline and Genomic Plasticity Underlying the Lifestyle Diversity in Rhizobiales. mSystems 5:e00438–20.

52. Garrido-Oter R, Nakano RT, Dombrowski N, Ma K-W, McHardy AC, Schulze-Lefert P. 2018. Modular Traits of the Rhizobiales Root Microbiota and Their Evolutionary Relationship with Symbiotic Rhizobia. Cell Host & Microbe 24:155-167.e5.

53. Carvalho FM, Souza RC, Barcellos FG, Hungria M, Vasconcelos ATR. 2010. Genomic and evolutionary comparisons of diazotrophic and pathogenic bacteria of the order Rhizobiales. BMC Microbiology 10:37.

54. Dziewit L, Baj J, Szuplewska M, Maj A, Tabin M, Czyzkowska A, Skrzypczyk G, Adamczuk M, Sitarek T, Stawinski P, Tudek A, Wanasz K, Wardal E, Piechucka E, Bartosik D. 2012. Insights into the Transposable Mobilome of Paracoccus spp. (Alphaproteobacteria). PLOS ONE 7:e32277.

55. Sun Y, Zhao J, Chen L, Liu Y, Zuo J. 2018. Methanogenic community structure in simultaneous methanogenesis and denitrification granular sludge. Frontiers of Environmental Science & Engineering 12:10.

